# Rab8, Rab11, and Rab35 coordinate lumen and cilia formation during Zebrafish Left-Right Organizer development

**DOI:** 10.1101/2022.10.31.514532

**Authors:** Abrar A. Aljiboury, Eric Ingram, Nikhila Krishnan, Favour Ononiwu, Debadrita Pal, Julie Manikas, Christopher Taveras, Nicole A. Hall, Jonah Da Silva, Judy Freshour, Heidi Hehnly

**Author notes:** Lead contact, correspondence, Twitter: @LovelessRadio. Authors contributed equally to this work and are listed in alphabetical order (Abrar A. Aljiboury, Eric Ingram, Nikhila Krishnan, Favour Ononiwu, Debadrita Pal).

## Abstract

An essential process during *Danio rerio’s* left-right organizer (Kupffer’s Vesicle, KV) development is for the majority of developing KV cells to form a motile cilium that extend into the KV lumen. Left-right beating of motile cilia within the KV lumen directs fluid flow to establishment the embryo’s left-right axis. However, when KV cells start to form cilia and how cilia formation is coordinated with KV lumen formation has not been examined. We identified that nascent KV cells form cilia at their centrosomes at random intracellular positions that then move towards a forming apical membrane containing cystic fibrosis transmembrane conductance regulator (CFTR). Using optogenetic clustering approaches, we found that Rab35 positive membranes recruit Rab11 to modulate CFTR delivery to the apical membrane, which is required for lumen opening, and subsequent cilia extension into the lumenal cavity. Once the intracellular cilia reach the CFTR positive apical membrane, Arl13b-positive cilia extend and elongate in a Rab8 dependent manner into the forming lumen once the lumen reaches an area of 300 μm^2^. These studies demonstrate the need to acutely coordinate Rab8, Rab11, and Rab35-mediated membrane trafficking events to ensure appropriate timing in lumen and cilia formation during KV development.

## INTRODUCTION

A fundamental question in cell biology is how a cilium is made during tissue formation. A primary or motile cilium is a microtubule-based structure that extends from the surface of a cell and can sense extracellular cues to transmit to the cell body. Defects in cilia formation can lead to numerous disease states collectively known as ciliopathies [1,2]. Foundational studies identified two distinct pathways for ciliogenesis *in vivo* using tissues from chicks and rats [3]. One mechanism for ciliogenesis which we refer to as extracellular, was found in lung cells where the centrosome first docks to the plasma membrane followed by growth of the ciliary axoneme into the extracellular space [4]. The second mechanism, which we refer to as intracellular, was identified in smooth muscle cells and fibroblasts where the centrosome forms a cilia first within a ciliary vesicle in the cell cytosol before docking to the plasma membrane [3]. These studies raise the possibility that different ciliated tissues construct their cilia differentially due to the nature of how a tissue develops. This presents an important hypothesis that variations in cilia formation mechanisms may occur *in vivo* during specific types of tissue morphogenesis.

Here we examine cilia formation during *Danio rerio* (zebrafish) organ of asymmetry (Kupffer’s Vesicle, KV) development. The KV is required to place visceral and abdominal organs with respect to the two main body axes of the animal [5]. KV formation begins from a sub-population of endoderm cells. The endoderm is induced by high levels of Nodal signaling during early development that contributes to the formation of the liver, pancreas, intestine, stomach, pharynx, and swim bladder [6]. A subset of the endoderm, called dorsal forerunner cells (DFCs) are precursors of the KV [7–9]. The number of DFCs range from 10-50 cells per embryo that can expand into >100 cells that make up the fully functional KV [10,11]. Early studies reported that these DFCs present as mesenchymal-like and are migratory. They lack clear apical/basal polarity (lack of aPKC distribution) until KV cells establish into rosette-like structures [9]. Apical polarity establishment of aPKC, at least in part, coincides with cystic fibrosis transmembrane conductance regulator (CFTR) accumulation at apical sites, which is a requirement for ultimate lumen expansion [9,12]. KV cell rosette-like structures can either form as multiple cells congressing to make a single rosette or cells assembling multiple rosettes which then transition to a single rosette-like structure. The rosette center is the site where a fluid-filled lumen forms and KV cells will then extend their cilia into [12]. Once KV cilia are formed they beat in a leftward motion to direct fluid flow essential for the establishment of the embryo’s left-right axis [13]. While much is known about KV post-lumen formation [5,14–17], little is known about the spatial and temporal mechanisms that regulate cilia formation during KV development.

*In vitro* cell culture assays have been used to identify regulators of lumen establishment or ciliogenesis and have identified 3 potential Rab GTPases that are involved in both: Rab11, Rab8, and Rab35 [18–24]. Here we investigated the role of these three Rab GTPases in KV development. Using a combination of depletion and optogenetic clustering approaches we have identified conserved yet unique roles for Rab8, Rab11 and Rab35 in coordinating KV lumen and cilia formation. While much is known about Rab8, Rab11, and Rab35, in ciliogenesis and/or lumen formation in the context of mammalian cell culture conditions, our findings were surprising in that Rab8 did not seem to affect lumen or cilia formation to a similar extent that it does in mammalian cell culture [18–21]. In mammalian cells, Rab8 and Rab11 work together in a GTPase cascade that is required for both cilia and lumen formation. However, in KV Rab35 and Rab11 seem to be coordinated and Rab8 is dispensable for lumen formation suggesting that specific cell types during potentially different developmental processes may have different dependencies on Rab GTPases that can be identified using developmental model systems such as zebrafish.

## RESULTS

Previous work in mammalian culture systems and preliminary morpholino studies in zebrafish KV have implicated Rab8, Rab11, and Rab35 in cilia and/or lumen establishment [18,19,21–24]. However, their cellular distribution during KV development has not been investigated, nor has it been positioned in relation to KV cilia formation. Foundational studies have demonstrated that at least one of the paralogs of Rab8, Rab11, and Rab35 is broadly expressed throughout zebrafish development, including KV [22,25–27]. In consideration of these findings, we assessed Rab8a, Rab11a, and Rab35 distribution in the zebrafish KV marked by the plasma membrane marker GFP-CAAX by expressing fluorescently tagged mRNA through injection (Figure 1A-B, S1A-C, Video S1) or in an endogenously GFP tagged transgenic line of Rab11 (Figure 1C, 1E, S1B [28]). Different stages of KV development, pre-rosette during the 1 somite stage (SS, 8-9 hours post fertilization, hpf), rosette during the 3 SS (10 hpf), and lumen during 6 SS (12 hpf, Figure 1A) were monitored using live or fixed embryo imaging preparations. We identified that Rab8 and Rab11 were broadly recruited to the apical membrane during rosette formation and remained there during lumen opening (Figure 1B-C, S1A). Embryos were fixed at the KV rosette stage (Figure 1E, top) and lumen stage (Figure 1E, bottom), and cilia were immunostained for acetylated tubulin. At the KV rosette stage cilia are organized intracellularly surrounded by Rab11 membranes organized at the center of the rosette, while Rab8 is organized at the base of the cilia where the centrosome resides (Figure 1E, centrosome staining in S1D). As the KV develops to a lumen stage, Rab11 reorganizes to the base with Rab8 (Figure 1E, S1B). Throughout the developmental stages Rab35 organized to cell boundaries (Figure 1D, S1C) with no specific localization to the cilia itself (Figure S1C).

**Figure 1.**
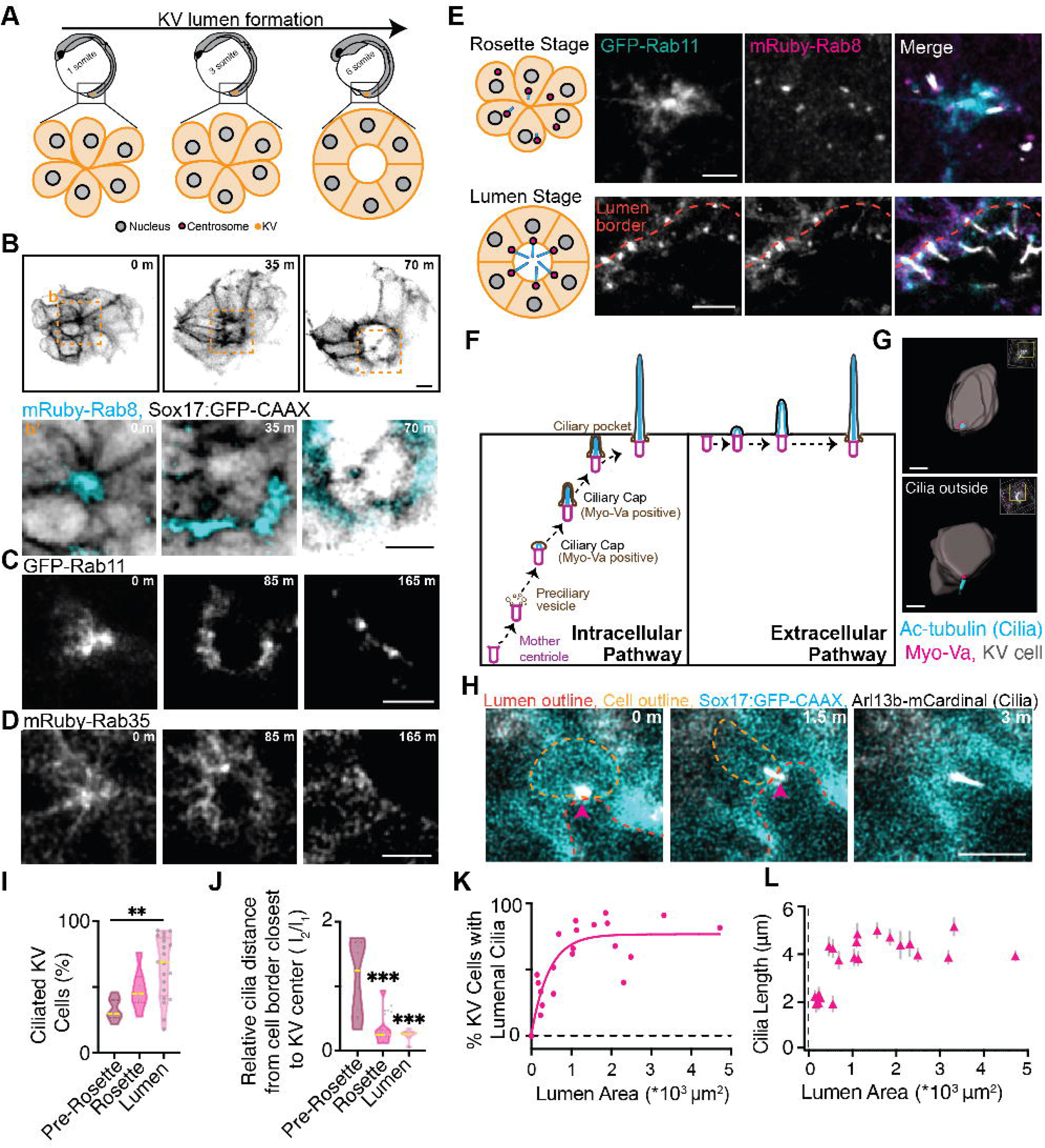
KV cilia form prior to KV lumen formation using an intracellular pathway. (**A**) Model depicting KV lumen formation across developmental stages of the zebrafish embryo. (**B-D**) Live confocal videos of mRuby-Rab8 (cyan, **B**), GFP-Rab11 (gray, **C**), and mRuby-Rab35 (gray, **D**) localization in KV cells marked by GFP-CAAX (inverted LUT, **B**) during lumen formation. Scale bar, 10μm. Refer to Video S1. (**E**) Left, model of KV developmental stages, rosette (top) and lumen (bottom), with centrosome (magenta) and cilia (cyan) positioning. Right, confocal micrographs with GFP-Rab11 (cyan), mRuby-Rab8 (magenta), and cilia (acetylated-tubulin, gray), shown. Bar, 10 μm. (**F)** Model demonstrating intracellular versus extracellular pathways for cilia formation. (**G**) 3D surface rendering of representative KV cells with cilia (acetylated-tubulin, cyan) inside versus outside of KV plasma membranes (KV membranes, Sox17:GFP-CAAX, gray), Myo-Va (magenta). Bar, 5 μm. **(H)** KV cell building and extending a cilium (Arl13b-mCardinal, inverted gray) into the lumen of the KV. KV plasma membranes (Sox17:GFP-CAAX) shown. Bar, 5 μm. (**I**) Percentage of ciliated KV cells at the different KV developmental stages. (**J**) Relative distance of cilia from cell border closest to KV center. (**I-J**) Shown as a violin plot with median (yellow line). One way ANOVA across KV developmental stages, n>7 embryos, **p<0.01. (**K**) Scatter plot demonstrating the percentage of KV cells with lumenal cilia per embryo in relation to KV lumen area. n=29 embryos. Goodness of fit R^2^= 0.8577. Please refer to Table S1 for additional statistical information. (**L**) Scatter plot depicting average cilia length within KV cells per embryo across n=29 embryos in relation to lumen area. Error bars, ± SEM.

A striking finding was that a significant population of KV cells started to form cilia at the centrosome in the cell body before a KV lumen formed (Figure 1E, S1D-E). During the pre-rosette stage, 33.25±3.33% of KV cells already had cilia; that increased to 48.06±5.94% at the rosette stage and averaged at 64.82±5.27% early on during lumen formation (Figure 1I, S1D-E). These studies suggested that KV cells were forming cilia before they had an extracellular space (KV Lumen) to position into and that KV lumen formation correlated with a significant increase in KV cells having cilia. Our findings suggest a cellular mechanism where cilia first are formed through an intracellular pathway that then will extend into the lumen (Figure 1F). To further test that cilia were forming through a potential intracellular pathway versus an extracellular pathway, volumetric projections of surface rendered KV cells were performed at the pre-rosette, rosette, and lumen stages. KV cell outlines were obtained using GFP-CAAX and cilia were immunostained using acetylated tubulin (Figure S1E) along with a marker for the ciliary membrane cap, Myosin Va (Myo-Va, [29,30], Figure 1F-G). Surface rendering using IMARIS software allowed for the spatial positioning of cilia in KV cells across KV developmental stages to be assessed (Figure S1E, Video S2). The boundaries of the cell (GFP-CAAX), cilia (acetylated tubulin), and Myo-Va were highlighted to create a three-dimensional space filling model of both cell, cilia, and ciliary cap (Figure 1G). We identified that as KV develops from pre-rosette to rosette, then to the lumenal stage, intracellular cilia surrounded by Myo-Va approach the apical membrane (Figure 1F-H, S1E, Video S2). Once a lumen is formed, the cilia extend into the developing KV lumen (Figure S1E, Video S2). Once the cilium extended out into the lumen, Myo-Va remained at the cilium’s base (Figure 1G). To identify if KV cell cilia were positioning towards the center of the KV cellular mass over the course of its development, we calculated the relative distance of cilia from the cell boundary closest to the KV center from the embryos shown in Figure S1D (modeled in Figure S1F; calculations in Figure 1J). When values approach 0, cilia are approaching the cell boundary closest to the KV center. This occurs significantly as KV cells transitioned from a pre-rosette organization to KV cells organized around a fluid filled lumen (Figure 1J). This suggests that KV cell cilia are constructed intracellularly, then positioned to the cell boundary closest to KV center where they are primed to extend their cilia into the forming lumen. These studies suggest a mechanism where KV cell cilia are forming through an intracellular pathway that recruits pre-ciliary vesicles positive for Myo-Va. These Myo-Va vesicles then form a ciliary cap for the cilia to grow within. The cilia with associated cap can then fuse with the plasma membrane and KV cilia can extend into the lumen (Figure 1F, left).

We next tested when cilia extend into the forming KV lumen. To do this we employed two strategies. We first imaged live Sox17:GFP-CAAX embryos that ectopically expressed the cilia marker Arl13b-mCardinal (Figure 1H). With the second strategy we fixed GFP-CAAX embryos at various lumen sizes ranging from 0 to 5*10^3^ μm^2^ and measured the percentage of KV cells that had lumenal cilia (Figure 1K) and cilia length (Figure 1L). These approaches demonstrated that cilia dock at the apical membrane during early lumen formation and then extend into the lumen (Figure 1H) once the lumen area approaches approximately 300 μm^2^ (Figure 1K). We then compared these studies to when cilia start to elongate (Figure 1L). We find that cilia, when inside a KV cell, can reach a length of 2.5 μm, but once a lumen is formed (300 μm^2^ in area), the cilia can extend into the lumen and grow to their final approximate length of 4 μm (Figure 1L).

To test the requirement of Rab11, Rab8, and Rab35 for KV cell cilia formation we employed two strategies, acute Rab GTPase optogenetic clustering assay (modeled in Figure 2A-B, S2A) and morpholino (MO) transcript depletion using MOs that have been previously characterized ([18,22,31], Figure S2B). Since Rab11, Rab8, and Rab35 are broadly expressed during zebrafish embryo development [22,25–27], we chose to employ an optogenetic strategy to acutely inhibit their function during KV development. This optogenetic strategy causes an acute inhibition of CIB1-Rab11-, CIB1-Rab8-, and CIB1-Rab35-associated membranes through a hetero-interaction between cyptochrome2 (CRY2) and CIB1 upon exposure to blue light during KV developmental stages [32–34]. Previous studies identified that upon blue light exposure, CIB1-Rab5 or CIB1-Rab11-associated membrane compartments cluster together creating an intracellular traffic jam and inhibiting the specific Rab’s membrane compartment from sorting intracellular cargo and regulating cellular functions [32–34]. Our studies herein find that optogenetically clustering Rab11- or Rab35-membranes during early KV development caused significant defects at 6 SS when the KV should have a lumen and many of the cells should be ciliated. Under control conditions (CRY2 injected) and Rab8 clustered conditions, 78.03±3.86% and 64.43±3.85% of KV cells formed cilia respectively, whereas Rab11 and Rab35 clustered embryos had a significant decrease in the percentage of ciliated cells (35.81±8.79% for Rab11, 49.72±5.50% for Rab35, Figure 2B-C, S2A). KV cells that could form cilia under Rab11- and Rab35-clustered conditions had most of their cilia stuck in the cell volume (Figure 2B, 2D). Rab11-, Rab8-, and Rab35-clustered cells that made cilia demonstrated significantly decreased cilia length (2.92±0.14 μm for Rab11, 3.15±0.08 μm for Rab8, and 2.02±0.05 for Rab35) compared to control CRY2 conditions (4.13±0.06 μm, Figure 2E). This significant decrease in cilia length with Rab11, Rab8, and Rab35 clustering, is consistent with Rab11, Rab8, or Rab35 depletion using morpholinos (Figure S2B-C). Interestingly, Rab8 clustering did cause a significant decrease in cilia length (Figure 2E), but not in the formation of cilia or its extension into the KV lumen (Figure 2C-D, S2A). These findings suggest that centrosomes that construct a cilium under Rab11- and Rab35-clustering conditions are unable to extend the cilium into the lumen and that this could be the underlying reason for cilia being significantly shorter in length. To test the role of Rab35-, Rab11- and Rab8-membranes in intracellular cilia positioning during KV development, the associated centrosome distances from the plasma membrane closest to KV center were measured under clustered conditions and compared to control conditions (CRY2). If centrosomes are positioning towards the KV center, then the number should approach 0. Rab11- and Rab35-clustered embryos measurements averaged around 0.70±0.10 and 0.75±0.11 respectively, whereas with Rab8 clustered and control conditions the centrosome distance approached 0 with a value of 0.23±0.06 and 0.30±0.03 (Figure 2F). These studies suggest that Rab11 and Rab35 coordinate centrosome and cilia positioning during KV development.

**Figure 2.**
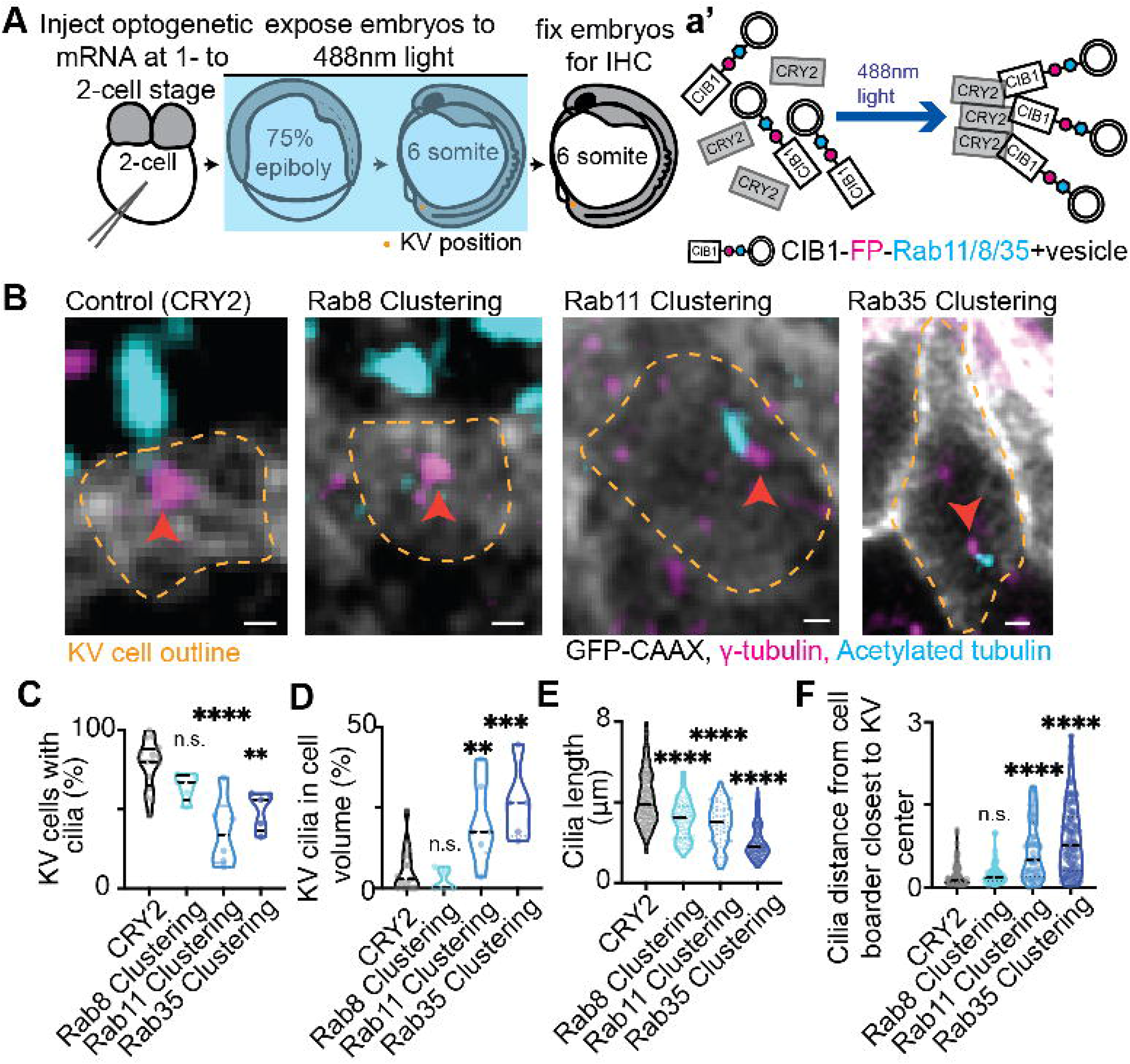
Cilia extension into the KV lumen requires Rab11- and Rab35-associated membranes, but not Rab8. (**A**) A model depicting the use of optogenetics to acutely block Rab-associated trafficking events during KV developmental stages. **(B**) Confocal micrographs of cilia (acetylated tubulin, cyan) in CRY2 (control), Rab8-, Rab11-, and Rab35-clustered Sox17:GFP-CAAX embryos (gray). Centrosomes denoted by γ-tubulin (magenta). Clusters not shown. Yellow dashed lines, KV cell cortical membranes. Orange arrow, centrosome. Bar, 2μm. (**C-F**) Violin plots of percentage of KV cells with cilia (**C**), percentage of KV cilia in cell volume (**D**), cilia length (**E**), and the relative distance of cilia from the cell boarder closest to KV center (**F**). One way ANOVA with Dunnett’s multiple comparison to CRY2 (control) was performed. n>4 embryos, n.s. not significant, **p<0.01, ***p<0.001, ****p<0.0001. Statistical results detailed in Table S1.

Our initial studies demonstrated that KV cilia extend into the lumen once the lumen reaches an area 300 μm^2^ (Figure 1K). Then KV cilia can reach their maximum length of approximately 4 μm (Figure 1L). These findings suggested that mechanisms regulating lumen formation may also play an important role in coordinating cilia formation. We tested the requirement of Rab11, Rab8, and Rab35 on KV lumen establishment using MO transcript depletion (Figure S2B, S3A-B) and the optogenetic clustering strategy (Figure 2A, 3A-C). With acute optogenetic clustering of Rab11- and Rab35-associated membranes in CFTR-GFP (Figure 3A) or Sox17:GFP-CAAX embryos (Figure 3B, S3C-D) we identified severe defects in KV lumen development that was consistent when depleting transcripts using MOs (Figure S2B, S3A-B) when comparing to control conditions (CRY2, Figure S3C; control MO, S3A-B). This was measured both by following lumen formation live using an automated fluorescent stereoscope set up for a set time frame (Figure 3B, S3C, Video S3) and at a fixed developmental endpoint (6 SS, Figure 3C, S3D). For live embryo analysis, Sox17:GFP-CAAX embryos were imaged just past 75% epiboly for over 4 hours, during this time, the Rab35 and Rab11 clustered embryos were not able to form a lumen when compared to control (CRY2) or Rab8-clustered embryos (Figure 3B, S3C, Video S3). When assessing at a fixed developmental endpoint (6SS, 12 hpf), we found that Rab11 and Rab35 clustered embryos presented with defects in forming a rosette (23.7% of embryos for Rab11, 21.3% for Rab35) or transitioning from a multiple rosette state to a single rosette state (18.3% of embryos for Rab11, 14.9% for Rab35, Figure 3C) compared to CRY2 embryos or Rab8 clustered embryos (98.8% and 98.6% form lumen, Figure 3C, S3C, Video S3). With Rab35 and Rab11 clustering, less than 50% of KVs were able to form a lumen (Figure 3C), and the lumens they did form were significantly decreased in size (Figure S3D). Interestingly, Rab35-clustered embryos were able to form separate rosettes that were not in the same cellular KV mass and in some cases one of the rosettes could transition to a small KV structure with a lumen (refer to rosette 1 in Figure 3A, quantification in Figure 3C, and additional example in Figure S3E). While a Rab11-Rab8 GTPase cascade during lumen formation has been proposed in the context of mammalian cell culture conditions [21], our findings were surprising in that acute Rab8 clustering conditions or Rab8 depletion conditions by MO does not affect lumen formation during KV development, but instead Rab11 and Rab35 play a predominant role.

**Figure 3.**
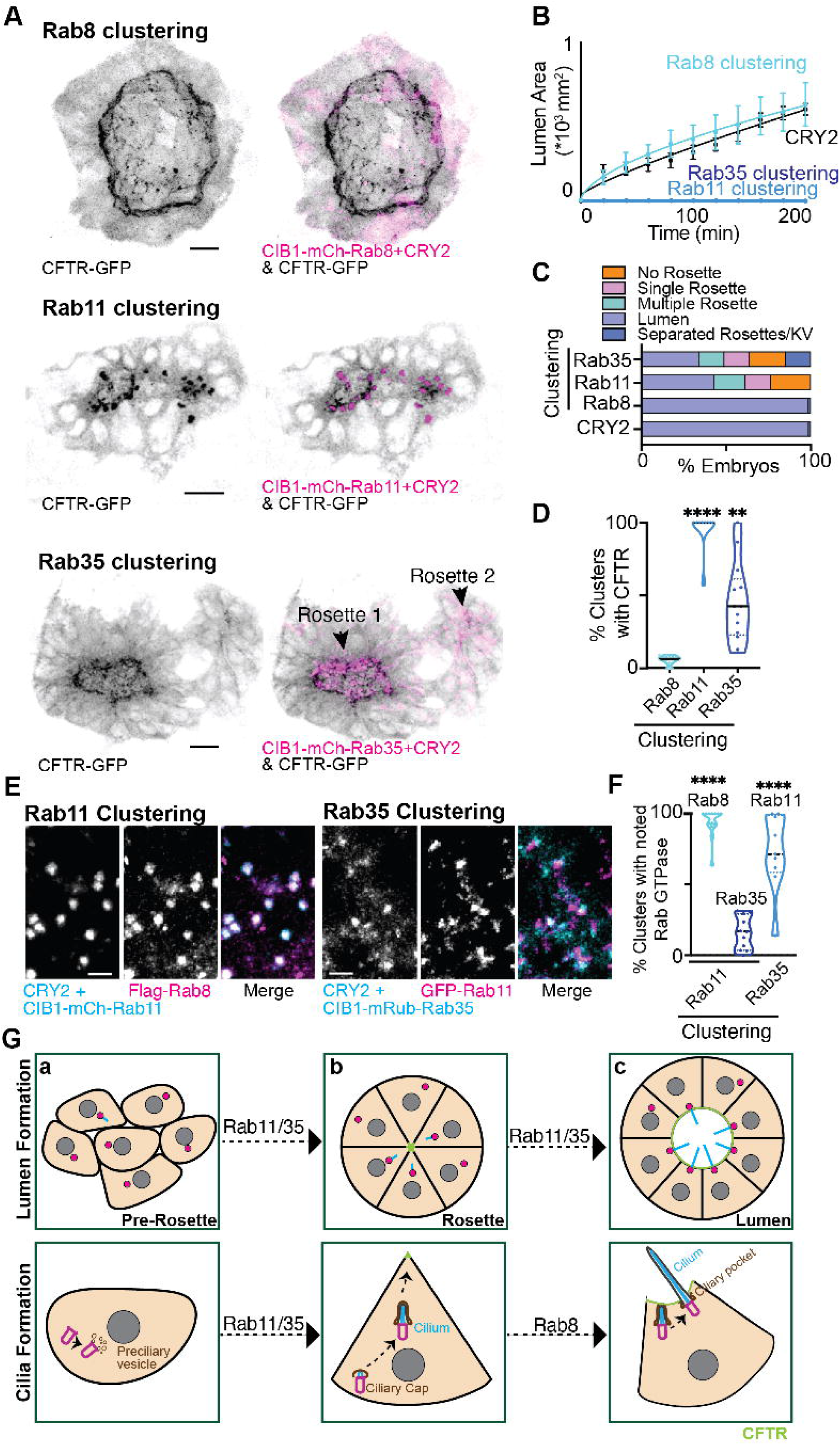
Rab11 and Rab35, but not Rab8, regulates KV lumen formation by mediating CFTR trafficking to the apical membrane. (**A)** Optogenetic clustering of Rab11, Rab8, and Rab35 (magenta) in KV cells. Localization with CFTR-GFP (inverted LUT) is shown. Bar, 20 μm. **(B)** KV lumen area over time (±SEM for n=3 embryos per condition) in control (CRY2 injection) and Rab8, Rab11 and Rab35 clustering conditions. See Figure S3C and Video S3 for representatives. **(C)** KV morphologies measured from optogenetically-clustered then fixed embryos at 12 SS (12 hpf). n>47 embryos per condition measured across n>9 clutches. (**D**) Percent of optogenetic clusters that colocalize with CFTR. n>9 embryos, **p<0.01, ****p<0.0001. (**E**) Optogenetic clustering of Rab11 and Rab35 (cyan). Rab11 clusters localization with Flag-Rab8 (magenta) or mRuby-Rab35 clusters with GFP-Rab11 (magenta) shown. Bar, 7 μm. (**F**) Percent of optogenetic clusters that colocalize with Rab8, Rab35, or Rab11 was calculated. n>9 embryos, ****p<0.0001. Statistical results detailed in Table S1. **(G)** Model depicting KV lumen and cilia formation across KV developmental stages. Centrosome depicted in magenta, cilia in cyan and CFTR in green. In short, a proportion of centrosomes start to assemble cilia at the pre-rosette stage that then reposition towards the center of the KV at the rosette stage in a Rab11 and Rab35 dependent manner. At this stage, Rab11 and Rab35 mediate CFTR transport to the apical membrane. The rosette stage then transitions to a lumen stage where most of the centrosomes can then locate at the CFTR-positive apical membrane and extend their cilia into the lumen where cilia can elongate to their full length in a Rab8 dependent manner.

Since both Rab11 and Rab35 optogenetic clustering, but not Rab8, resulted in lumen formation defects we wanted to examine whether they disrupted CFTR recruitment to the apical membrane. CFTR is a master regulator of fluid secretion into lumenal spaces. CFTR is transported through the secretory pathway to the apical membrane where it mediates chloride ion transport from inside the cell to outside the cell. Loss of CFTR-mediated fluid secretion impairs KV lumen expansion leading to laterality defects [12]. Our studies find that Rab11 optogenetic clustering causes a severe defect in CFTR delivery to the apical membrane where CFTR-GFP becomes trapped in Rab11- and Rab35-clustered membrane compartments (Figure 3A, 3D). With both Rab11 and Rab35 clustering, there was significantly less CFTR that was able to be delivered to forming apical membranes. This is consistent with defects in KV rosette and lumen formation observed with Rab11 and Rab35 clustered embryos (Figure 3C). Interestingly, some Rab35 clustered embryos assemble multiple rosettes in a KV, with one rosette being competent for lumen formation but defective in expansion (Figure 3A). When this occurs, we find that the rosette that is competent in opening has some CFTR localized to the apical membrane (Rosette 1, Figure 3A), as opposed to the secondary rosette that cannot open (Rosette 2, Figure 3A). No defect in CFTR delivery to the apical membrane was noted with Rab8 optogenetic clustering (Figure 3A, 3D), consistent with the lack of observed defects in lumen formation with both optogenetic clustering (Figure 3B-C) and depletion of Rab8 using morpholinos (Figure S3A-B).

Some GTPases are known to work together on the same membrane compartment. For instance, Rab11 and Rab8 were reported to function together in a GTPase cascade on recycling endosomes to regulate cellular events such as lumen formation and ciliogenesis. In this situation, Rab11 acts upstream of Rab8 by recruiting the Guanine Exchange Factor (GEF) for Rab8, Rabin8 [18,19,21]. Based on our findings that both Rab11 and Rab35 cause defects in lumen formation and CFTR trafficking, we asked if Rab11, Rab35, and/or Rab8 could act on the same membrane compartment. To test this, we performed optogenetic clustering of Rab35 or Rab11 and determined whether clustering one recruited Rab11, Rab35, or Rab8. Optogenetic clustering of Rab11 resulted in the recruitment of Rab8 but not Rab35 (Figure 3E-F, S3F). This is consistent with the idea that a Rab11 cascade may still exist between Rab11 and Rab8, but that this cascade is not required for CFTR transport or KV lumen formation. It also suggests that Rab11 is not acting upstream of Rab35. Interestingly, upon optogenetic clustering Rab35 membranes, Rab11 becomes co-localized (Figure 3E-F) suggesting that Rab35 is upstream of Rab11. In summary, we find that Rab35 may act upstream of Rab11 to ensure appropriate lumen formation through managing CFTR trafficking to the forming apical membrane.

Mechanistically we have found that during the KV pre-rosette stage, cells start to assemble a cilium inside the cell (Figure 3Ga). The cilium and associated centrosome are repositioned inside the cell towards the center of the KV cell mass at a similar time KV cells are rearranging into a rosette like structure (Figure 3Gb). This movement of the centrosome and rearranging into a rosette like structure also rely on Rab11 and Rab35 (Figure 3Gc). Forming and expanding the lumen likely depends on the ability of Rab11 and Rab35 to mediate CFTR transport to the apical membrane (Figure 3Gc, top). Once this occurs, the KV cell cilia can extend and elongate into the lumen in a Rab8 dependent manner (Figure 3Gc, bottom).

The consequences associated with KV lumen expansion and cilia formation can have downstream developmental defects that include defects in the left-right development of the brain, heart, and gut [5]. Based on this, we wanted to examine the developmental defects associated with acute optogenetic clustering of Rab8-, Rab11-, or Rab35-associated membranes during KV development (Figure 4A). Embryos expressing CIB1-Rab8, -Rab11, or -Rab35 with CRY2 were exposed to blue light to induce clustering at 75% epiboly when KV precursor (Dorsal Forerunner) cells are first visualized until 6 SS (12 hpf) when KV lumen is forming. The interaction between CIB1 and CRY2 is dependent on blue light, and after blue light is removed membranes can become unclustered [32]. At 6 SS blue light was removed and the embryos were left to develop to high pec (42 hpf). Gross phenotypes observed with animals having Rab8-, Rab11-, or Rab35-optogenetically clustered membranes included a significant increase in animals with curved tails (Figure 4B-C). Rab35 clustering specifically resulted in a significant increase of animals displaying no tails and/or a one-eye phenotype when compared to control conditions (CRY2, Figure 4B-C). We then examined heart development due to its laterality being easily assessed in live embryos (Figure 4D). Using a cmlc2:GFP transgenic line to label zebrafish heart cells specifically, we assessed the process of heart looping. Abnormal heart looping includes reversed looping, no loop, or bilateral heart looping (Figure 4D-E, Video S4). Over 70% of animals presented with abnormal heart looping when Rab8-, Rab11-, or Rab35-was acutely clustered during KV developmental stages compared to control animals expressing CRY2 (13.55±1.59%). While this was not surprising for Rab11 and Rab35 optogenetic clustering during KV development due to embryos presenting with severe lumen and cilia formation defects (Figure 2, 3), this was surprising for Rab8 optogenetic clustering conditions where embryos formed normal lumens but had slightly shorter cilia (Figure 2D). This suggests that even subtle defects in cilia length and potential function during KV development may result in significant developmental defects. Taken together, these studies propose that Rab8-, Rab11-, and Rab35-mediated membrane trafficking is necessary for forming a functional KV during development.

**Figure 4.**
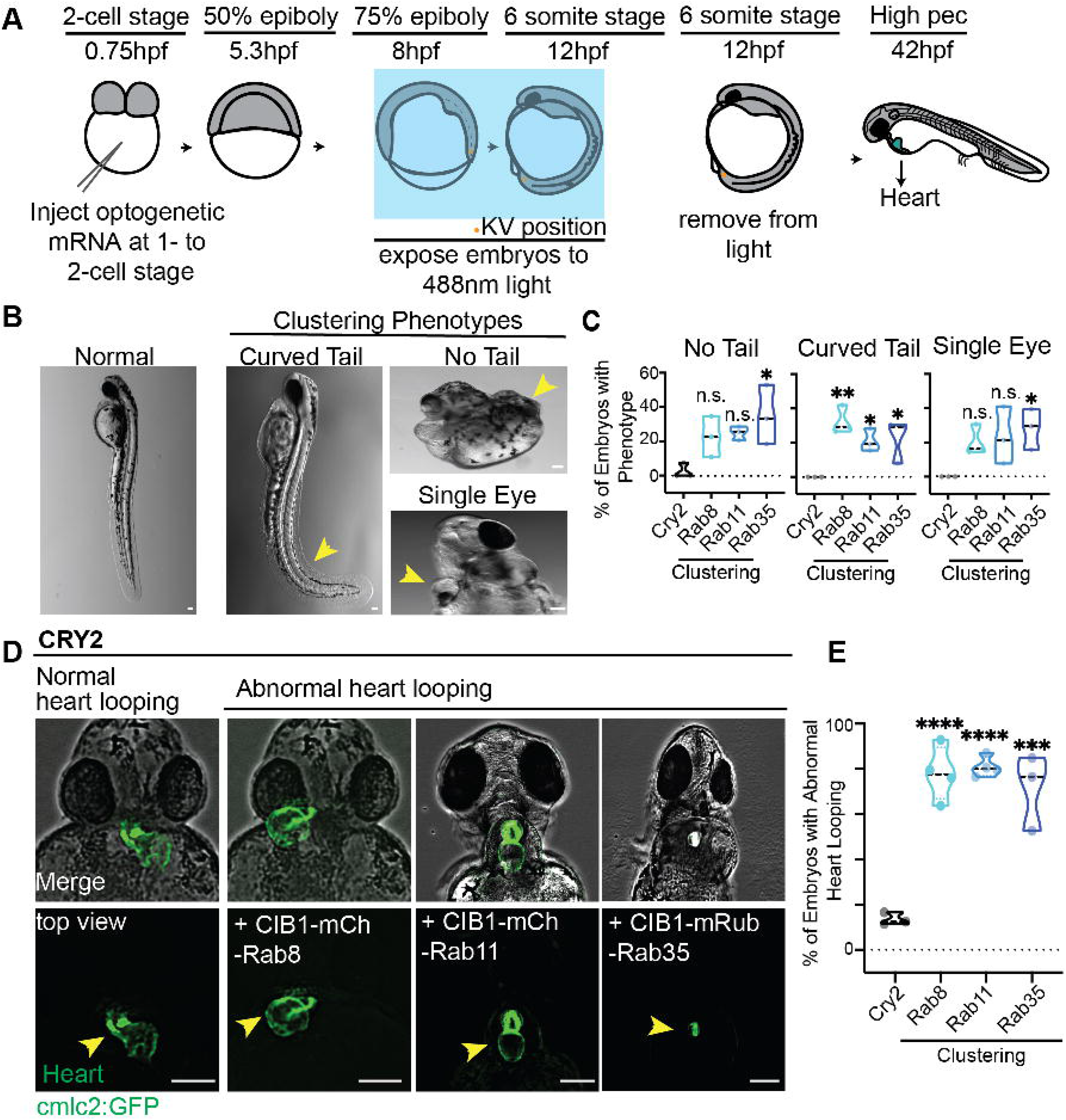
Acute optogenetic disruption of Rab8, Rab11, and Rab35 membranes during KV development results in left-right asymmetry defects. (**A**) A model depicting the use of optogenetics to acutely block Rab GTPase-associated trafficking events during KV developmental stages and assessment of downstream developmental consequences at 42 hpf. (**B**) Images demonstrate characterized developmental phenotypes observed that include curved tail, no tail, and single eye. Yellow arrows point to abnormalities. Bar, 100 μm. (**C**) Violin plot displaying percentage of embryos displaying a no tail, curved tail, or single eye phenotype (shown in (**B**)) over n>3 clutches across the optogenetic clustering conditions compared to control. *p<0.05, and **p<0.01. (**D**) Images demonstrate characterized abnormal heart looping in clustered embryos compared to normal leftward heart looping in control CRY2 embryos. Refer to Video S4. Bar, 100 μm. (**E**) Violin plot displaying percentage of embryos with abnormal heart looping (shown in (**D**)) n>3 clutches across the optogenetic clustering conditions compared to control. ***p<0.001 and ****p<0.0001. Statistical results detailed in Table S1.

## DISCUSSION

While select Rab GTPases have been extensively studied, most of them have not been assigned a detailed function or localization pattern during early embryonic vertebrate development. Rab GTPases have approximately 60 genes in vertebrates, with each Rab GTPase localizing to specific intracellular membrane compartments in their GTP-bound (active) form. These active Rabs then bind to effector proteins to aid in various steps in membrane trafficking some of which will facilitate cilia, polarity, and/or lumen formation [35,36]. Because Rab GTPases are potentially required for a variety of cellular functions and developmental contexts, we needed to employ a strategy to acutely disrupt their function. Herein, we used an optogenetic strategy that takes advantage of Rab GTPases membrane association, where the Rab GTPase of interest is attached to CIB1 and we express CIB1’s optogenetic binding partner CRY2. Upon exposure to blue light, CIB1 will form heteromeric complexes with CRY2 essentially causing the Rab associated membranes to cluster together and become non-functional (modeled in Figure 2A, 4A). This approach is versatile in developmental models due to its acute triggering and reversibility. For instance, we can acutely cluster Rab-associated membranes during a specific developmental stage, and then release the clustering through the removal of blue-light and examine downstream developmental consequences (Figure 4A). Zebrafish embryos are an ideal developmental system for this work due to their optical transparency and external development making them easily accessible to blue light addition [33,34]. In these studies we focused on three Rab GTPases (Rab8, Rab11, and Rab35) that have been linked to lumen and cilia formation in mammalian cell culture models [18,19,21,24] and employed a combination of optogenetic approaches and traditional depletion approaches using MO to examine their roles *in vivo* during left-right organizer development.

The left-right organizer is a conserved tissue in vertebrate embryos that establishes the embryo’s left-right axis. We used zebrafish as a model system to better understand left-right organizer development. In zebrafish, previous foundational studies identified that cells that make up the left-right organizer (KV) need to assemble into a cyst like structure surrounding a fluid filled lumen, with the majority of KV cells having a motile cilium [9,17,37]. Motile cilia beating in a left-right manner within the KV lumen directs fluid flow, which is essential for the establishment of the embryo’s left-right axis [5]. However, when KV cells start to form cilia and how cilia formation is coordinated with KV lumen formation had yet to be identified. Our studies have established that KV precursor cells, DFCs, assemble a cilium inside the cell before the KV cells start to assemble into a cyst-like structure (Figure 1E, S1D-E, Video S2). We noted that cilium and the associated centrosome reposition inside the cell towards the center of the KV cellular mass at a similar time KV cells are rearranging into a rosette like structure (Figure 1J, S1D-E), a pre-requisite structure that precedes lumen formation. This movement of the centrosome and rearranging into a rosette like structure relies on the small GTPases Rab11 and Rab35 (Figure 2 and 3). Specifically, we find that Rab35 acts upstream of Rab11 on the same membrane compartment, likely recycling endosomes [38], to assist in forming and expanding the lumen by regulating the delivery of CFTR to the apical membrane (Figure 3). Previous foundational work identified that CFTR recruitment to the apical membrane is a requirement for lumen expansion [12]. We identified that once the lumen expands to a specific area (Figure 1F, K), which is mediated by Rab11 and Rab35 (Figure 3), the KV cilia can extend and elongate into the lumen in a Rab8 dependent manner (Figure 2).

Interestingly, the only significant defect we identified with acute disruption of Rab8 was cilia length (Figure 2E), whereas with Rab11 and Rab35 we found defects in KV development that included rosette formation and transition to forming a lumen (Figure 3A-C, S3A-E), along with a defect in cilia formation (Figure 2, S2). This was surprising due to previous reports identifying a GTPase cascade between Rab11 and Rab8 that was needed for lumen formation and for cilia formation in mammalian tissue culture [18,19,21,39,40]. While we argue that this cascade may not be required for lumen or cilia formation in KV cells, it may still be intact in regulation of cilia length (Figure 2E, S2C). Our findings demonstrate that both conserved and divergent mechanisms are likely involved in cilia formation dependent on the developmental requirements of the tissue being formed. For instance, there may be a possible connection between Rab35 and Rab11 that is coordinated during cilia and lumen formation, where both Rab35 and Rab11 clustered membranes result in the sequestration of CFTR (Figure 2A, 2D), and that Rab35 clustering results in the partial recruitment of Rab11 (Figure 2E-F). Interestingly, there is no colocalization with Rab35 and cilia (Figure S1C), unlike Rab11 (Figure 1E). One potential unique mechanistic possibility that we have already touched upon is that Rab35 and Rab11 work together in coordinating lumen formation through CFTR transport (Figure 3G). In support of this scenario, Rab11 or Rab35 clustering prevents CFTR from accumulating appropriately at the apical membrane, resulting in incomplete lumen formation that would consequently cause cilia to remain inside the cell. This is indeed the case where we find with acute inhibition of Rab11 and Rab35 associated membrane compartments, a significant increase in KV cells have internalized cilia compared to control and Rab8 membrane inhibition (Figure 2D). These same conditions cause defects in lumen formation (Figure 3A-C, S3A-E) and the KVs that do form a lumen are significantly decreased in area and rarely reach that 300 μm^2^ lumen area threshold that is required for cilia to extend into the lumen (Figure 1H, 1K-L). An additional more conserved mechanism for Rab11, like what is reported in mammalian cell culture, is a direct role at the cilium where Rab11 localizes to (Figure 1E). In this scenario, Rab11 can regulate cilia formation and potential elongation in a cascade with Rab8. We argue that this cascade is likely in place based on our findings that acute inhibition of Rab11-associated membranes through optogenetic clustering recruits Rab8 to these membranes, but clustering Rab8 does not recruit Rab11. These findings suggest that Rab11 is upstream of Rab8 and can recruit Rab8 to the same membrane compartment to potentially regulate cilia elongation (Figure 2E, modeled in Figure 3G).

Our findings demonstrate that both conserved and divergent mechanisms for cilia formation likely exist, and Rab GTPases relative roles are likely dependent on the developmental requirements of the tissue being formed. Our studies validate zebrafish to be a versatile model to identify the potential mechanisms of function for Rab GTPases *in vivo*.

## ACKNOWLEDGEMENTS

We thank the Michel Bagnat lab at Duke University School of Medicine for sharing their eGFP-Rab11 transgenic zebrafish lines. This work was supported by National Institutes of Health grants R01GM127621 (H.H.) and R01GM130874 (H.H.).

## AUTHOR CONTRIBUTIONS

A.A., D.P., J.S., H.H., N.K., J.M., C.T., E.I., N.A.H and F.O. designed, performed, and analyzed experiments; H.H. wrote manuscript; J.F. provided molecular reagents and zebrafish husbandry. All authors provided edits. H.H. oversaw project.

## FIGURE LEGENDS

**Figure S1.**
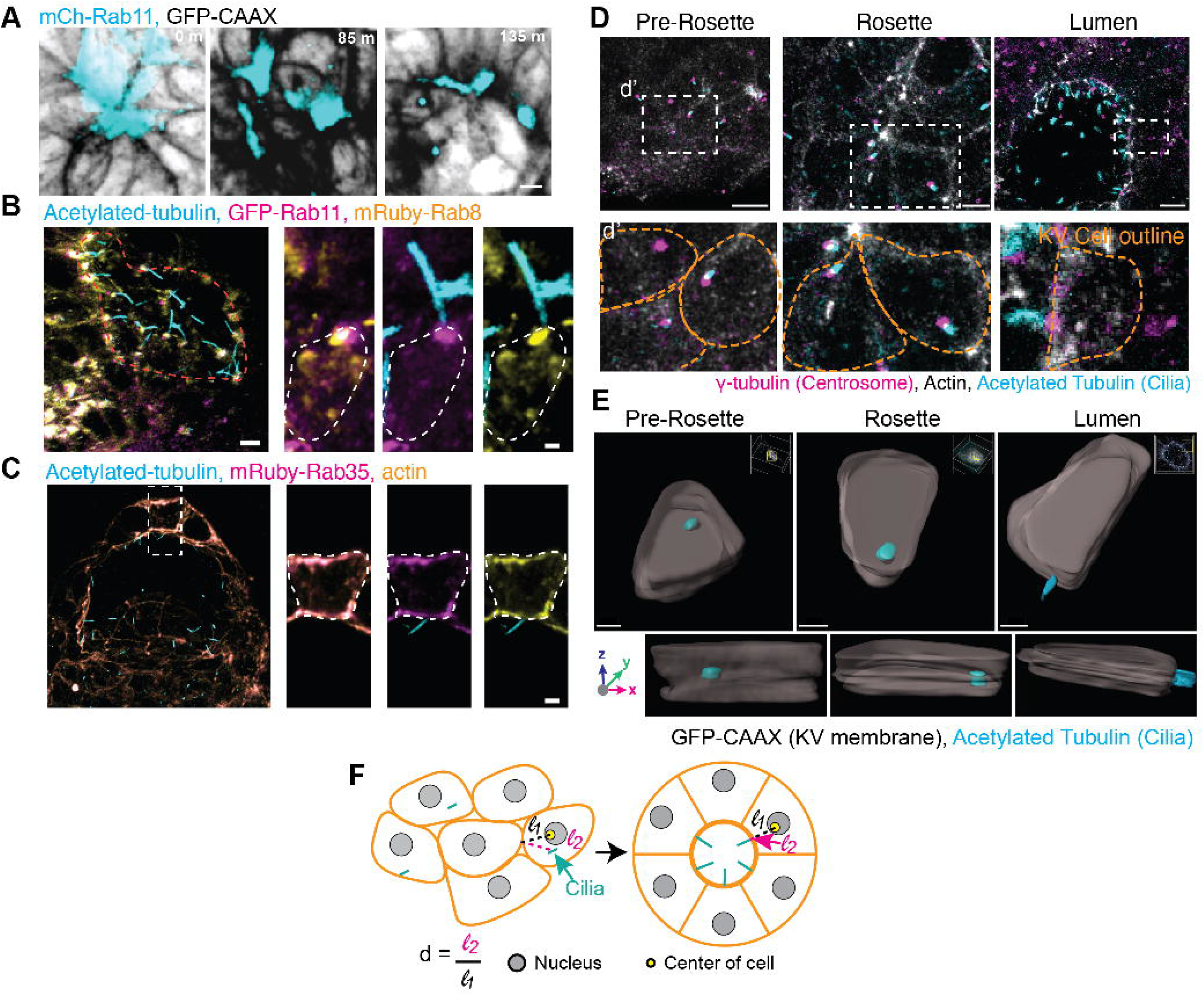
KV cilia form prior to KV lumen formation using an intracellular pathway. (**A**) Live confocal videos of mCherry-Rab11 (cyan) localization in KV cells during lumen formation. KV plasma membrane noted with GFP-CAAX (inverted gray). Scale bar, 10 μm. **(B-C)** Confocal micrographs of KV lumen stage with cilia (acetylated-tubulin, cyan), GFP-Rab11 (magenta, **B**), mRuby-Rab8 (yellow, **B**), mRuby-Rab35 (magenta, **C**), and actin (yellow, **C**). Bar, 2 μm. **(D)** Confocal micrographs of KV developmental stages with cilia (acetylated-tubulin, cyan), centrosome (γ-tubulin, magenta), and actin (phalloidin, gray). Scale bar, 10 μm. (**d’**) Magnified insets from (**D**) depicting centrosome and cilia positioning in KV cells at different KV developmental stages. Bar, 7 μm. (**E**) 3D surface rendering of a representative KV cell during pre-rosette, rosette, and lumen KV developmental stages with cilia (acetylated-tubulin, cyan) and KV plasma membranes (KV membranes, Sox17:GFP-CAAX, gray) rendered. Refer to Video S2. Bar, 5 μm. **(F)** Model depicting quantification of relative distance of the cilium from the cell border closest to KV center. Cilia, cyan. Nucleus, gray. Center of KV cells, yellow. Pink dashed line is distance of cilium from cell membrane. Black dashed line is distance of cell center to cell membrane.

**Video S1**. *Videos of Rab8, Rab11, and Rab35 distribution during KV lumen formation*. Live confocal videos of mRuby-Rab8 (cyan), GFP-Rab11 (gray), and mRuby-Rab35 (gray) localization in KV cells during lumen formation. KV plasma membranes (Sox17:GFP-CAAX) shown with actin and Rab8 (inverted gray). Bar, 10 μm. Refer to Figure 1B-D.

**Video S2**. *Cilia cellular positioning in 3D*. 3D surface rendering from Figure S1A of a single KV cell at the KV pre-rosette, rosette, or lumen stage rotated 360 degrees around the X-axis. Bar, 5 μm. Inset shows full KV with cilia (cyan) and KV plasma membrane (Sox17:GFP-CAAX, gray). Refer to Figure S1E.

**Figure S2.**
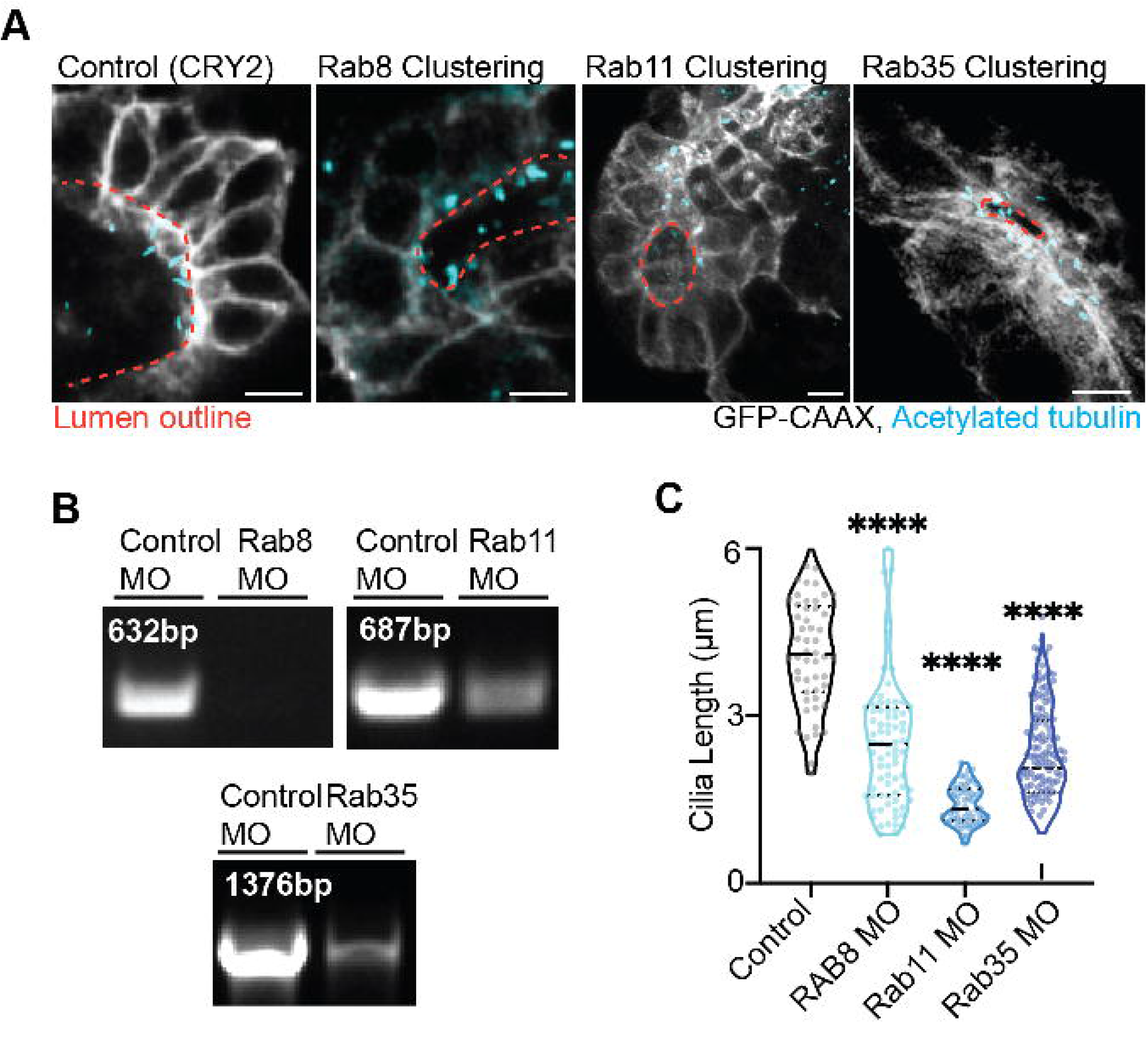
Cilia extension into the KV lumen requires Rab11- and Rab35-associated membranes, but not Rab8. (**A**) Confocal micrographs of cilia (acetylated tubulin, cyan) in CRY2 (control), Rab8-, Rab11-, and Rab35-clustered Sox17:GFP-CAAX embryos (gray). Clusters not shown. Lumen outline is orange dashed lines. Bar, 10 μm. **(B)** Agarose gel demonstrating RT-PCR of Rab8, Rab11, and Rab35 MO treatment compared to control MO conditions. Amplification of Rab8, Rab11, and Rab35 transcripts shown. NC, negative control. (**C**) Violin plot depicting cilia length from control, Rab8, Rab11, and Rab35 MO treatment. Dots represent individual cilia length values. Median denoted by line. One-way ANOVA with Dunnett’s multiple comparison test, compared to CRY2. ****p<0.0001. Statistical results detailed in Table S1.

**Figure S3.**
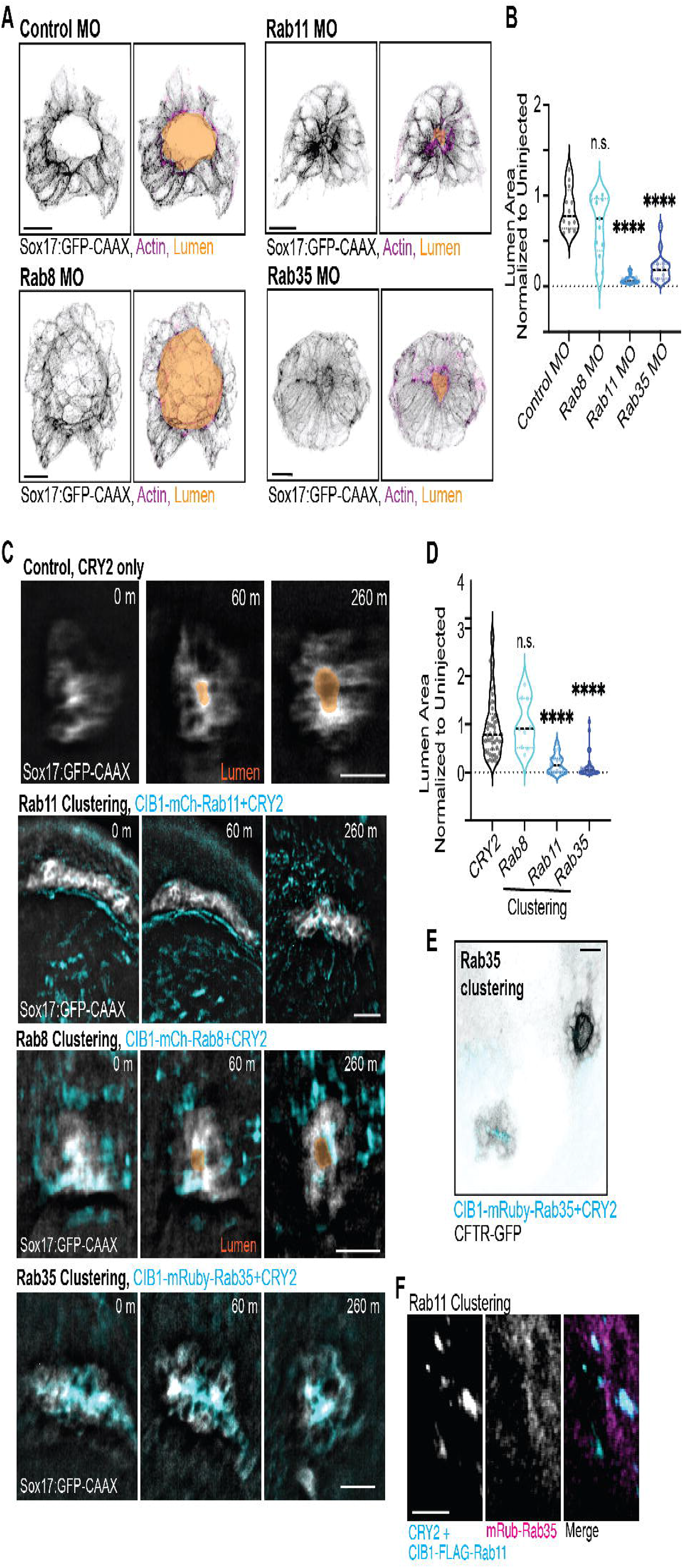
Rab11 and Rab35, but not Rab8, regulates KV lumen formation by mediating CFTR trafficking to the apical membrane. **(A)** Representative 3D rendering of KV under Rab8, Rab11, and Rab35 MO treatment. Lumen trace (orange), cell membrane (GFP-CAAX, inverted LUT), and actin (magenta) shown. Bar, 25 μm. **(B)** Violin plot depicting lumen area normalized to uninjected control values in control, Rab8, Rab11 and Rab35 MO injected embryos. Dots represent individual KV values. Median denoted by line. One-way ANOVA with Dunnett’s multiple comparison test, compared to control MO. n>12 embryos, n.s. not significant, ****p<0.0001. **(C)** Optogenetic clustering of Rab11 and Rab35 blocks KV lumen formation compared to CRY2 control and Rab8. Imaged on an automated fluorescent stereoscope. Bar, 50 μm. KV marked with Sox17:GFP-CAAX, lumens highlighted in orange, clusters shown in cyan. Refer to Video S3 and Figure 3B. **(D)** Violin plot depicting lumen area from Rab8, Rab11, and Rab35 clustering conditions normalized to uninjected control values. Dots represent individual KV values. Median denoted by line. One-way ANOVA with Dunnett’s multiple comparison test, compared to CRY2. n>9 embryos, ****p<0.0001. **(E)** Representative image of optogenetic clustering of Rab35 (cyan) in KV cells; CFTR-GFP (inverted LUT) shown. Bar, 25 μm. **(F)** Optogenetic clustering of Rab11 in KV cells. Rab11 clusters localization with mRuby-Rab35 (magenta) shown. Bar, 7 μm. Statistical results detailed in Table S1.

**Video S3**. *Rab11 and Rab35 modulate KV lumen formation*. Optogenetic clustering of Rab11 and Rab35 blocks KV lumen formation compared to Rab8. Embryos imaged on automated fluorescent stereoscope every 10 min. Bar, 100 μm. KV marked with Sox17:GFP-CAAX. Refer to Figure 3B and S3C.

**Video S4**. *Acute optogenetic disruption of Rab8, Rab11, and Rab35 membranes during KV development results in abnormal heart looping*. Stereo microscope video showing ventral view of a 48 hpf cmlc2:GFP (green) fish marking the heart. In CRY2 control, heart tube loops to the left. Rab8-, Rab11-, and Rab35-optogenetic clustered zebrafish reveal defective heart loop phenotype from Figure 4D. Size bar, 100 μm. 0.25s time interval.

## EXPERIMENTAL PROCEDURES

### Resource Availability

#### Lead contact

For further information or to request resources/reagents, contact Lead Contact, Dr. Heidi Hehnly

#### Materials availability

New materials generated for this study are available for distribution.

#### Data and code availability

All data sets analyzed for this study are displayed.

### Experimental model and subject details

#### Fish Lines

Zebrafish lines were maintained using standard procedures approved by Syracuse University IACUC (Institutional Animal Care Committee) (Protocol #18-006). Embryos were raised at 28.5°C and staged (as described in [41]). Wildtype and/or transgenic zebrafish lines used for live imaging and immunohistochemistry are listed in key resource table (Table S2).

#### Method Details

##### Antibodies

Antibody catalog information used in mammalian cell culture and zebrafish embryos are detailed in key resource table (Table S2).

##### Plasmids and mRNA

Plasmids were generated using Gibson cloning methods (NEBuilder HiFi DNA assembly Cloning Kit) and maxi-prepped before injection and/or transfection. mRNA was made using mMESSAGE mMACHINE™SP6 transcription kit. See key resource table for a list of plasmid constructs and mRNA used.

##### Morpholinos

Morpholinos (MO) were ordered from Gene Tools. Previously characterized Rab8, Rab11, and Rab35 MO sequences were used from [18,22,40]. See Supplementary key resource table in Table S2 for a list of morpholinos used.

##### RNA extraction and RT-PCR

Total RNA was extracted from either an isolated embryo or several embryos injected with control, Rab8, Rab11 or Rab35 morpholinos using TRIzol reagent. The RT-PCR was performed on each sample using OneTaq One-Step RT-PCR Kit (see key resource table) with the forward primers “tcagtatggcgaagacctacgat”, “gttagcatggctactgcctaatcac”, “gtaatgagcgactgactgctgac” and reverse primers “tcttcacagtagcacacagcga”, “catgtcattgtctcggcggtc”, “gtgcaaggagaaaaataagatcaagttagagaatca” for Rab8, Rab11 and Rab35 consecutively. RT-PCR reaction was run using the following cycling conditions: 48 °C for 30 min, 94 °C for 1min followed by 40 cycles of 94 °C for 15 sec, 54 °C (Rab8 and Rab11) or 53 °C (Rab35) for 30 sec, 68 °C for 2 minutes with final extension at 68 °C for 5 min.

##### Immunofluorescence

Fluorescent transgenic and/or mRNA injected embryos (refer to strains and mRNAs in key resource table, and for injection protocols refer to [42,43]) were staged at Kupffer’s Vesicle (KV) developmental stages as described in [33,44] and fixed using 4% paraformaldehyde with 0.1% triton-100. Standard immunofluorescent protocols were carried out (refer to [43]). Embryos were then embedded in low-melting 2% agarose (see key resource table) with the KV positioned at the bottom of a #1.5 glass bottom MatTek plate (see key resource table) and imaged using the spinning disk confocal microscope or laser scanning confocal microscope (see details below).

##### Imaging

Zebrafish embryos were imaged using Leica DMi8 (Leica, Bannockburn, IL) equipped with a X-light V2 Confocal unit spinning disk equipped with a Visitron VisiFRAP-DC photokinetics unit, a Leica SP8 (Leica, Bannockburn, IL) laser scanner confocal microscope (LSCM) and/or a Zeiss LSM 980 (Carl Zeiss, Germany) with Airyscan 2 confocal microscope. The Leica DMi8 is equipped with a Lumencore SPECTRA X (Lumencore, Beaverton, OR), Photometrics Prime-95B sCMOS Camera, and 89 North-LDi laser launch. VisiView software was used to acquire images. Optics used with this unit are HC PL APO x40/1.10W CORR CS2 0.65 water immersion objective, HC PL APO x40/0.95 NA CORR dry and HCX PL APO x63/1.40-0.06 NA oil objective. The SP8 laser scanning confocal microscope is equipped with HC PL APO 20x/0.75 IMM CORR CS2 objective, HC PL APO 40x/1.10 W CORR CS2 0.65 water objective and HC PL APO x63/1.3 Glyc CORR CS2 glycerol objective. LAS-X software was used to acquire images. The Zeiss LSM 980 is equipped with a T-PMT, GaASP detector, MA-PMT, Airyscan 2 multiplex with 4Y and 8Y. Optics used with this unit are PL APO x63/1.4 NA oil DIC. Zeiss Zen 3.2 was used to acquire the images. A Leica M165 FC stereomicroscope equipped with DFC 9000 GT sCMOS camera was used for staging and phenotypic analysis of zebrafish embryos.

##### Optogenetic experiments in zebrafish embryos

Tg(sox17:GFP-CAAX), Tg*BAC*(cftr-GFP), Tg(sox17:GFP), Tg(sox17:DsRed) and TgKIeGFP-Rab11a zebrafish embryos were injected with 50-100 pg of CRY2 and/or CIB1-mCherry-Rab11, CIB1-mCherry-Rab8 or CIB1-mRuby-Rab35 at the one cell to 4 cell stage. Embryos were allowed to develop in the dark until uninjected embryos reached the 75% epiboly stage where we can screen embryos for KV cells and expose them to 488nm light using the NIGHTSEA fluorescence system until the six-somite stage [33]. Embryos were then fixed and immunostained (refer to [43]).

##### Analysis of Zebrafish developmental defects and heart looping defects following acute optogenetic clustering

Zebrafish embryos injected with optogenetic constructs were exposed to 488nm light from 8 hpf-12 hpf as described in [33]. Embryos were incubated at 28.5°C until 42 hpf. Zebrafish were manually dechorionated using forceps and mounted in 2% agarose before imaging. Heart loop assessment and imaging were carried out on Leica M165 FC stereomicroscope equipped with DFC 9000 GT sCMOS29camera. A Plan Apochromat 1X objective and GFP excitation emission filter was used. Images were acquired using LAS-X software and post-image processing was done using thunder imaging system from Leica. Lateral view and ventral view of zebrafish were obtained from bright field imaging. Time lapse video of heart looping was performed at 0.25 seconds interval. Heart looping was characterized by leftward, rightward, and severely defective looping. Gross embryo phenotypes were categorized into no tail, curved tail, and single eye phenotypes. Categorization was performed over 787 embryos over n>3 clutches with at least 88-274 embryos per condition.

##### Image and data analysis

Images were processed using FIJI/ImageJ. Graphs and statistical analysis were produced using Prism 9 software. Surface rendering (refer to [33]) and analysis of KV cells were performed using Bitplane IMARIS software. Videos were created using FIJI/ImageJ or IMARIS. Cilia length was measured as the distance from the base of the cilia to the tip using line function in IMARIS. For percentage of ciliated KV cells, the number of cells with cilia was counted and represented as a percentage over the total number of cells in the cyst forming tissue.

##### Relative cilia distance from cell border closest to KV center

the distance from cilia to the cell membrane closest to KV center (l2) was measured and divided by the distance of the center of the cell (nucleus) to the cell’s membrane closes to KV center(l1); d=l2/l1 (refer to Figure S1F). This was done for KV cells with positive cilia staining at each developmental KV stage.

##### Calculating colocalization of CFTR and Rab GTPases with select optogenetic clusters

From fixed embryos the total number of Rab GTPase clusters were counted for each KV. The number of Rab clusters that had CFTR or Rab GTPase being tested overlapping with the Rab GTPase cluster was counted and presented as a percentage.

##### Statistical Analysis

Unpaired two-tailed t-tests and one way ANOVA were performed using PRISM9 software. **** denotes a p-value<0.0001, *** p-value<0.001, **p-value<0.01, *p-value<0.05, n.s. not significant. For further information on detailed statistical analysis see supplemental table 1.

