## Supplemental Table S1 for "Rab8, Rab11, and Rab35 coordinate lumen and cilia formation during Zebrafish Left-Right Organizer development"

**Table S1. Detailed statistical analysis of results reported in this study.**

| Figure | Category | n cell | n embryo | n Clutch | Statistical Test | Parameters | Result | p-value |
|---|---|---|---|---|---|---|---|---|
| 1I | Pre-Rosette | N/A | n=7 | n>3 | One Way ANOVA | F (2,31) = 6.917 | ** | 0.0033 |
| | Rosette | N/A | n=7 | | | | | |
| | Lumen | N/A | n=20 | | | | | |
| 1J | Pre-Rosette | n=8 | n=1 | N/A | One Way ANOVA | F (2,21) = 13.62 | N/A | N/A |
| | Rosette | n=8 | n=1 | | | | *** | 0.0006 |
| | Lumen | n=8 | n=1 | | | | *** | 0.0003 |
| 1K | KV cells with lumenal cilia across lumen area | N/A | n=29 | n>3 | N/A | N/A | N/A | N/A |
| 1L | N/A | N/A | n=29 | n>3 | N/A | N/A | N/A | N/A |
| 2C | CRY2 | N/A | n=16 | n>2 | One Way ANOVA | F (3, 31) = 12.27 | N/A | N/A |
| | Rab8 clustering | N/A | n=16 | | | | n.s. | 0.2732 |
| | Rab11 clustering | N/A | n=5 | | | | **** | <0.0001 |
| | Rab35 clustering | N/A | n=15 | | | | ** | 0.0009 |
| 2D | CRY2 | N/A | n=16 | n>2 | One Way ANOVA | F (3, 31) = 13.27 | N/A | N/A |
| | Rab8 clustering | N/A | n=5 | | | | n.s. | 0.8825 |
| | Rab11 clustering | N/A | n=6 | | | | ** | 0.0070 |
| | Rab35 clustering | N/A | n=5 | | | | **** | <0.0001 |
| 2E | CRY2 | n=433 | n=17 | n>3 | One Way ANOVA | F (3, 874) = 189.2 | N/A | N/A |
| | Rab8 clustering | n=145 | n=4 | | | | **** | <0.0001 |
| | Rab11 clustering | n=61 | n=5 | | | | **** | <0.0001 |
| | Rab35 clustering | n=239 | n=8 | | | | **** | <0.0001 |
| 2F | CRY2 | n=91 | n=6 | n=3 | One Way ANOVA | F (3, 299) = 46.89 | N/A | N/A |
| | Rab8 clustering | n=69 | n=6 | | | | n.s. | 0.9545 |
| | Rab11 clustering | n=49 | n=4 | | | | **** | <0.0001 |
| | Rab35 clustering | n=94 | n=9 | | | | **** | <0.0001 |
| S2C | Control MO | n=52 | n=3 | n>2 | One Way ANOVA | F (3, 319) = 74.45 | N/A | N/A |
| | Rab8 MO | n=105 | n=7 | | | | **** | <0.0001 |
| | Rab11 MO | n=55 | n=3 | | | | **** | <0.0001 |
| | Rab35 MO | n=111 | n=5 | | | | **** | <0.0001 |
| 3B | CRY2 | N/A | n=3 | n=3 | N/A | N/A | N/A | N/A |
| | Rab8 clustering | N/A | n=3 | | N/A | N/A | N/A | N/A |
| | Rab11 clustering | N/A | n=3 | | N/A | N/A | N/A | N/A |
| | Rab35 clustering | N/A | n=3 | | N/A | N/A | N/A | N/A |
| 3C | CRY2 | N/A | n=80 | n>9 | N/A | N/A | N/A | N/A |
| | Rab8 clustering | N/A | n=72 | | N/A | N/A | N/A | N/A |
| | Rab11 clustering | N/A | n=93 | | N/A | N/A | N/A | N/A |
| | Rab35 clustering | N/A | n=47 | | N/A | N/A | N/A | N/A |
| 3D | Rab8 Clustering | N/A | n=8 | n=3 | One Way ANOVA | F (2, 27) = 43.87 | N/A | N/A |
| | Rab11 Clustering | N/A | n=9 | | | | **** | <0.0001 |
| | Rab35 Clustering | N/A | n=13 | | | | ** | 0.0015 |
| 3F | Rab11 Clustering-Rab8 | N/A | n=13 | n>2 | One Way ANOVA | F (2, 30) = 63.95 | N/A | N/A |
| | Rab11 Clustering-Rab35 | N/A | n=9 | | | | **** | <0.0001 |
| | Rab35 Clustering-Rab11 | N/A | n=13 | | | | **** | <0.0001 |
| S3B | Control MO | N/A | n=12 | n=3 | One Way ANOVA | F (3, 58) = 51.52 | N/A | N/A |
| | Rab8 MO | N/A | n=13 | n=2 | | | n.s. | 0.0839 |
| | Rab11 MO | N/A | n=21 | n=2 | | | **** | <0.0001 |
| | Rab35 MO | N/A | n=16 | n=4 | | | **** | <0.0001 |
| S3D | CRY2 | N/A | n=50 | n=11 | One Way ANOVA | F (3, 94) = 18.43 | N/A | N/A |
| | Rab8 clustering | N/A | n=9 | n=3 | | | **** | <0.0001 |

| | | | | | | | | |
|---|---|---|---|---|---|---|---|---|
| | Rab11 clustering | N/A | n=16 | n=4 | | | n.s. | 0.9962 |
| | Rab35 clustering | N/A | n=23 | n=2 | | | **** | <0.0001 |
| 4C | No tail-CRY2 | N/A | n=85 | n=3 | One Way ANOVA | F (3, 8) = 4.645 | N/A | N/A |
| | No tail- Rab8 clustering | | n=135 | | | | n.s. | 0.1235 |
| | No tail- Rab11 clustering | | n=120 | | | | n.s. | 0.0860 |
| | No tail- Rab35 clustering | | n=88 | | | | * | 0.0167 |
| | Curved Tail-CR2 | N/A | n=85 | n=3 | One Way ANOVA | F (3, 8) = 8.210 | N/A | N/A |
| | Curved Tail- Rab8 clustering | | n=135 | | | | ** | 0.0034 |
| | Curved Tail- Rab11 clustering | | n=120 | | | | * | 0.0363 |
| | Curved Tail- Rab35 clustering | | n=88 | | | | * | 0.0254 |
| | Single Eye- CRY2 | N/A | n=85 | n=3 | One Way ANOVA | F (3, 8) = 3.707 | N/A | N/A |
| | Single Eye- Rab8 clustering | | n=135 | | | | n.s. | 0.1292 |
| | Single Eye- Rab11 clustering | | n=120 | | | | n.s. | 0.0800 |
| | Single Eye- Rab35 clustering | | n=88 | | | | * | 0.0367 |
| 4E | CRY2 | N/A | n=161 | n=3 | One Way ANOVA | F (3, 10) = 30.37 | N/A | N/A |
| | Rab8 Clustering | | n=264 | n=4 | | | **** | <0.0001 |
| | Rab11 Clustering | | n=269 | n=4 | | | **** | <0.0001 |
| | Rab35 Clustering | | n=88 | n=3 | | | *** | 0.0001 |
