## Supplemental Table S2 for "Rab8, Rab11, and Rab35 coordinate lumen and cilia formation during Zebrafish Left-Right Organizer development"

**Table S2. SUPPLEMENTARY KEY RESOURCE TABLE**

| Reagent or resource | Source | Identifier |
|---|---|---|
| **Antibodies** | | |
| Acetylated Tubulin | Sigma Aldrich | T6793: RRID: AB_477585 |
| Gamma-tubulin | Sigma Aldrich | T5192; RRID: AB_261690 |
| Anti-GFP (Chicken) | GeneTex | GTX13970: AB_371416 |
| Anti-GFP (Rabbit) | Molecular Probes | A-11122: AB_221569 |
| Anti-Flag (Mouse) | Sigma-Aldrich | F3165-.2MG |
| Anti-Flag (Rabbit) | Sigma | F7425 |
| Myosin 5a | Novus Biologicals | NBP1-92156 |
| Alexa Fluor Anti-Rabbit 488 | Life Technologies | A21206; RRID: AB_2535792 |
| Alexa Fluor Anti-Rabbit 568 | Life Technologies | A10042; RRID: AB_2534017 |
| Alexa Fluor Anti-Rabbit 647 | Life Technologies | A31573; RRID: AB_2536183 |
| DyLight 405-AffiniPure Donkey Anti-Mouse IgG (H+L) | Jackson ImmunoResearch | 715-475-150 |
| Alexa Fluor Anti-Mouse 488 | Life Technologies | A21202; RRID: AB_141607 |
| Alexa Fluor Anti-Mouse 568 | Life Technologies | A10037; RRID: AB_2534013 |
| Alexa Fluor Anti-Mouse 647 | Life Technologies | A31571; RRID: AB_162542 |
| **Chemicals, Peptides, and Recombinant Proteins** | | |
| DAPI | Sigma Aldrich | D9542-10mg |
| Alexa Fluor 647 Phalloidin | Cell Signaling Technology | 8940S |

| | | |
|---|---|---|
| Agarose | Thermo Fischer | 16520100 |
| BSA | Fisher Scientific | BP1600-100 |
| BIO BASIC Maxi Prep Kit | BIO BASIC | 9K-0060023 |
| Dimethylsulphoxide | Fisher Scientific | BP231-100 |
| Paraformaldehyde | Fisher Scientific | O4042-500 |
| Phosphate Buffered Saline | Fisher Scientific | 10010023 |
| Life Technologies Prolong Diamond Antifade mount with DAPI | Fisher Scientific | P36971 |
| 35 mm Dish\| No.1.5. coverslip\| 20 mm Glass Diameter | MatTek Corporation | P35G-1.5-20-C |
| Molecular Probes Prolong Gold Antifade mount | Fisher Scientific | P36934 |
| Triton X-100 | Fisher Scientific | BP151500 |
| Tween 20 | ThermoFischer | BP337500 |
| Sodium Chloride | Fisher Scientific | BP358 |

| | | |
|---|---|---|
| NEBuilder HiFi DNA assembly Cloning Kit | New England BioLabs | E5520S |
| mMESSAGE mMACHINETMSP6 | Invitrogen | AM1340 |
| OneTaq One-Step RT-PCR Kit | New England Biolabs | E5315S |
| **Experimental models, organisms, and strains** | | |
| Zebrafish | Zebrafish International Resource Center | AB-Wildtype |
| Zebrafish | Zebrafish International Resource Center | Tg (Sox17:DsRed) |
| Zebrafish | (Dasgupta and Amack, 2016) | Tg (sox17:GFP-CAAX)sny101 |
| Zebrafish | (Navis et al., 2013) | TgBAC(cftr-GFP) |
| Zebrafish | (Levic et al., 2020) | TgKIeGFP-Rab11a |
| Zebrafish | Zebrafish International Resource Center | Tg(sox17:GFP) |
| Zebrafish | Megason Lab | βactin:EMTB-3xGFP; cmlc2:GFP |
| **mRNA and Morpholinos** | | |
| CRY2 | (Rathbun et al., 2020) | Plasmid: pCS2-CRY2; Addgene Plasmid #140572 |
| CIB1-mCherry-Rab11a | (Rathbun et al., 2020) | Plasmid: pCS2-CIB1-mCherry-Rab11a; Addgene Plasmid #140573 |

| | | |
|---|---|---|
| CIB1-mCherry-Rab8a | This paper | Plasmid: pCS2-CIB1-mCherry-Rab8a |
| CIB1-mRuby-Rab35 | This paper | Plasmid: pCS2-CIB1-mRuby-Rab35 |
| FLAG-Rab8 | This paper | Plasmid: pCS2-FLAG-Rab8 |
| FLAG-Rab11 | This paper | Plasmid: pCS2-FLAG-Rab11 |
| mRuby-Rab8a | This paper | Plasmid: pCS2-mRuby-Rab8a |
| mCherry-Rab11 | (Krishnan *et al.*, 2022) | Plasmid: pCS2- mCherry-Ra11 |
| mRuby-Rab35 | This paper | Plasmid: pCS2- mRuby-Rab35 |
| Arl13b-mCardinal | This paper | Plasmid: pCS2- Arl13b-mCardinal |
| **Morpholinos** | | |
| Control MO | vivo standard control morpholinos | Gene Tools |
| Rab8 MO | (Omori *et al.*, 2008; Lu *et al.*, 2015) | GAAGACATAAATACCTATCGTCGAG |
| Rab11 MO | (Westlake *et al.*, 2011) | GTATTCGTCGTCTCGTGTCCCCAT |
| Rab35 MO | (Kuhns *et al.*, 2019) | TGCAGCTTCACGCCTCTCTCCAGCA |
| **Software and algorithms** | | |
| ImageJ/FIJI | NIH and Laboratory for Optical and Computational Instrumentation | https://imagej.net/Fiji |

| | | |
|---|---|---|
| IMARIS, Bitplane | Oxford Instruments | https://imaris.oxinst.com/ |
| PRISM9 | GraphPad | https://www.graphpad.com/scientific-software/prism/ |
| LAS-X Software | Leica Microsystems | https://www.leica-microsystems.com/products/microscope-software/p/leica-las-x-ls/ |
| VisiView | Visitron | https://www.visitron.de/products/visiviewr-software.html |
